# Benchmarking Imputation Methods for Single-Cell RNA Sequencing Data Using Peripheral Blood Mononuclear Cells from Acute Myocardial Infarction Patients

**DOI:** 10.64898/2026.08.23.746230

**Authors:** Prathiksha Ramesh, Maria Fyta

**Affiliations:** Department of Biological Sciences and Engineering, Indian Institute of Technology Gandhinagar, Gujarat, India; Computational Biotechnology, RWTH Aachen University, Worringerweg 3, 52074 Aachen, Germany; Center for Computational Life Sciences (CCLS), RWTH Aachen University, Pauwelstrasse 19, 52074 Aachen, Germany

**Keywords:** scRNA-sequencing, Machine Learning, dropout imputation, peripheral blood mononuclear cells, benchmarking, acute myocardial infarction, GAN

## Abstract

Acute myocardial infarction (AMI) remains one of the leading causes of mortality worldwide, and the following post-effects, such as post-AMI inflammation and tissue repair, involve peripheral blood mononuclear cells playing a critical role. The influence of imputation methods in biological data is assessed with respect to high-resolution single-cell RNA sequencing (scRNAseq) data relevant to these cells. Still scRNAseq data often encounter a lot of dropout events, leading to sparse and noisy datasets, hampering downstream results. To assess the influence of the missingness in the data, we artificially impose different levels of dropout in available scRNAseq data by leveraging various imputation techniques. Specifically, we introduce artificial missingness at 10%, 20%, and 30% levels under a missing completely at random (MCAR) framework, repeated across 10 independent runs. We benchmarked six imputation strategies - MAGIC, IterativeImputer, KNNImputer, Mean Imputation, SoftImpute, and a Generative adversarial network (GAN) - based approaches using multiple evaluation metrics: marker gene preservation, clustering consistency (Adjusted Rand Index - ARI), gene-wise correlation with ground truth, and structural separation (silhouette scores). The results clearly underline that no single imputation method dominated across all metrics. Overall, Mean and KNN imputers showed limited recovery across all benchmarks. GAN excelled in global transcriptional recovery and SoftImpute in preserving biologically meaningful cell-type signals. Our results highlight the importance of selecting the imputation methods as part of the pre-processing step towards the downstream biological questions related to transcriptome recovery, detection of marker genes, or maintaining cell-type-specific resolution.

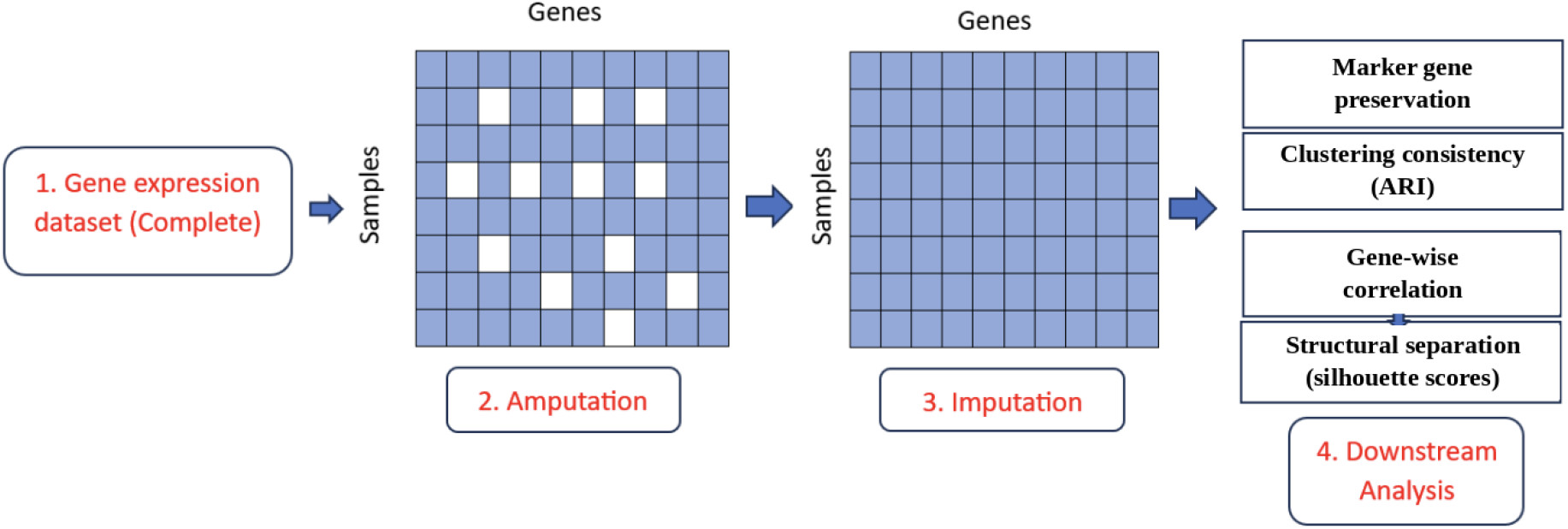

## Introduction

Acute Myocardial Infarction (AMI), or commonly known as a heart attack, is one of the leading causes of mortality worldwide, arising due to the permanent damage to the heart muscle [1]. Such AMIs can occur with atherosclerotic plaque rupture in a coronary artery [2]. Plaque rupture promotes thrombus formation by exposing thrombogenic components and thereby activating the clotting cascade. Beyond these mechanical events, during and after rupture, immunological PBMCs are involved in the process. Post AMI, PBMCs increase in number and migrate to the damaged heart tissue, initiating the inflammatory and healing processes [3]. Initially, flow cytometry, wherein fluorescent antibodies were used to label specific cell surface proteins and intracellular molecules, was used to analyze PBMCs after AMI to understand their inflammatory and reparative roles [4]. Later on, bulk RNA sequencing was used to identify changes in immune cell composition and gene expression profiles, providing insights into systemic immune response to heart damage. Both of those techniques are often validated and combined, like single-cell RNA sequencing, to provide a more comprehensive understanding of the cellular and molecular changes following AMI.

Single-cell RNA sequencing (scRNA-seq) [14] is a prominent tool used to study the immune responses and complex cellular changes occurring after acute myocardial infarction by providing a high-resolution view of gene expression in individual cells [6]. scRNA-seq helps understand cellular diversity and heterogeneity, characterizing cellular dynamics, identifying immune responses, discovering new therapeutic targets, fibroblast heterogeneity, endothelial cell states, immune cell signatures, and biomarker discovery [5]. Despite its transformative potential, scRNA-seq suffers from a fundamental technical challenge: dropout events. In scRNA-seq, a dropout event occurs when a gene that has been expressed in a cell is not detected by the sequencing method, resulting in a zero in the gene expression matrix. These events are the result of low mRNA abundance, technical inefficiencies, and stochastic gene expression. Such dropouts in scRNA sequencing data lead to data sparsity, noise, and inaccuracies, which potentially challenge downstream analyses, including cell clustering, differential gene expression analysis, and trajectory inference [8,9]. As a result, biological conclusions may be incomplete or biased if dropout is not properly addressed.

In order to mitigate the challenges posed by dropouts, various computational methods have been developed to “impute” or estimate the missing expression values. Such methods often leverage information from similar cells or genes to infer the missing data. While many of the imputation methods have been benchmarked in generic scRNA-seq contexts, their impact on disease-specific datasets, such as AMI PBMCs, has not been systematically studied [10]. In order to investigate systemic immune responses in AMI, a recent landmark study, “*Single Cell RNA Sequencing of peripheral blood mononuclear cells from acute myocardial infarction*,” accessed from NCBI GEO on 08 September 2025 [19] was used. This dataset comprises ∼ 82,550 single cells from 10 patients (split into plaque rupture vs non-rupture groups), annotated into 6 discrete immune clusters (monocytes, T / B / NK / progenitors). The dataset includes raw matrices, features, and barcode files generated using the 10x Genomics Chromium platform for peripheral blood mononuclear cells (PBMCs) from acute myocardial infarction (AMI) patients, categorized into plaque rupture and non-rupture groups. These raw matrices were used for downstream quality control, normalization, and imputation analyses. It thus captures the transcriptional diversity of circulating PBMCs during the acute phase of myocardial infarction. For computational benchmarking, this dataset serves as a ground truth reference due to its depth and quality.

In the present study, we have used this baseline expression profile and systematically simulated MCARs [7] of varying levels of artificial dropout (10%, 20%,30%) to mimic increasing degrees of sparsity. These simulated datasets were then subjected to multiple imputation methods, and outputs were compared to the original ground truth data to assess their ability to recover true gene expression patterns, clustering fidelity, and biologically meaningful gene-gene correlations. To date, to our knowledge, no systematic effort has been made to evaluate the effect of different imputation strategies, specifically in the context of PBMCs from AMI patients. Using the recently generated single-cell dataset of AMI as a reference or ground truth, our work aims to provide a focused benchmark highlighting how imputation influences biological interpretation in this disease setting. By doing so, we guide future cardiovascular research toward more accurate, reproducible, and biologically meaningful use of single-cell data.

## Methods

### Dataset and preprocessing

We used a single-cell RNA sequencing dataset from the NCBI Gene Expression Omnibus (GEO) database [19]. This dataset contains a total of 10 samples collected from AMI patients, of which five with and five without plaque rupture, and captures the transcriptional changes occurring in circulating immune cells. Raw count matrices were processed in Python (Scanpy v1.9.5) following standard scRNA-seq quality control (QC) (we have followed common workflows [20]. Low-quality cells with fewer than 200 detected genes or with greater than 5% mitochondrial gene expression were excluded. Genes expressed in fewer than three cells were also filtered. Following QC, the data were normalized using log-normalization, and highly variable genes were identified for downstream analysis.

### Simulation of missingness and imputation methods

In order to mimic dropout effects commonly observed in scRNA-seq data, we have introduced artificial sparsity by randomly masking a fraction of observed counts at three levels: 10%, 20%, and 30%, respectively. Each analysis was repeated across 10 independent runs to account for statistics, ensuring the robustness of the evaluation. We have systematically benchmarked a diverse set of imputation methods, including statistical, graph-based, and Machine Learning (ML) approaches, all of which were based on publicly available Python packages. These include:

a. **Markov Affinity-based Graph Imputation of Cells (MAGIC)**, a graph-based smoothie method [11].
b. **IterativeImputer**, which predicts missing values step by step using other genes [15].
c. **KNNImputer**, which fills gaps based on the most similar nearest cells [16,17].
d. **Mean Imputation**, a simple baseline that replaces missing values with the average [18].
e. **SoftImpute**, which uses a matrix completion approach [12].
f. **Generative Adversarial Networks (GANs)**, which use deep learning to learn patterns in the data [13].

In order to assess the performance of these learning models and each imputation strategy applied, we have computed four different metrics with a biological context:

- **Marker gene preservation:** we examined whether imputation methods retained expression of canonical immune cell markers relative to the ground truth reference dataset. Marker gene expression for each imputed dataset was compared against the corresponding ground truth values using correlation-based scores, restricted to markers present within the ground truth’s highly-variable-gene feature set (top 2,000 genes; see *Dataset and preprocessing*). This yielded eight evaluable markers spanning platelet (PF4, PPBP), B-cell (CD79A, MS4A1), NK/cytotoxic (GZMB, FGFBP2, KLRC1), and monocyte/granulocyte (S100A12) lineages.
- **Clustering consistency:** we assessed the effect of imputation on cell clustering using the Adjusted Rand Index (ARI). Following dimensionality reduction and clustering, ARI was calculated by comparing the imputed clusters to those from the unaltered ground truth dataset. This metric directly tests whether imputation distorts or preserves cell identity assignment.
- **Gene-wise correlation:** To measure how well imputation restored overall expression patterns, we computed Pearson correlations for each gene between the imputed and ground truth datasets. These values were averaged across all genes to obtain global gene recovery performance.
- **Structural separation:** Finally, we used Silhouette scores to quantify the degree of cluster separation in low-dimensional embeddings. While this indicates structural clarity, we interpreted these results with caution, as over-smoothing may inflate silhouette values without reflecting genuine biological recovery.

## Results & Discussion

In the following, we briefly elaborate on the main features and most important insights gained from our analysis. We gather the statistics across 10 independent masking computations for each of the imputation schemes mentioned and the three missing fractions (10%, 20%, and 30%) each. We go through each of the metrics analyzed and focus on the most impactful outcome. In view of the marker gene preservation in order to evaluate the ability of imputation methods to retain biologically meaningful signals, we assessed the recovery of canonical immune cell markers against the ground truth dataset. The heatmap in Figure 1 shows the eight markers common to both the ground truth reference and all imputed datasets, spanning platelet, B-cell, NK/cytotoxic, and monocyte lineages. The results represented in the figure clearly reveal that SoftImpute, MAGIC, and GAN consistently preserved marker gene expression patterns, with average preservation scores ranging from 0.75 to 0.84. On the other hand, Mean and KNN imputers showed a substantial loss of signal, with some marker scores approaching zero. The Iterative approach produced intermediate results, with performance varying across markers and dropout levels.

**Figure 1:**
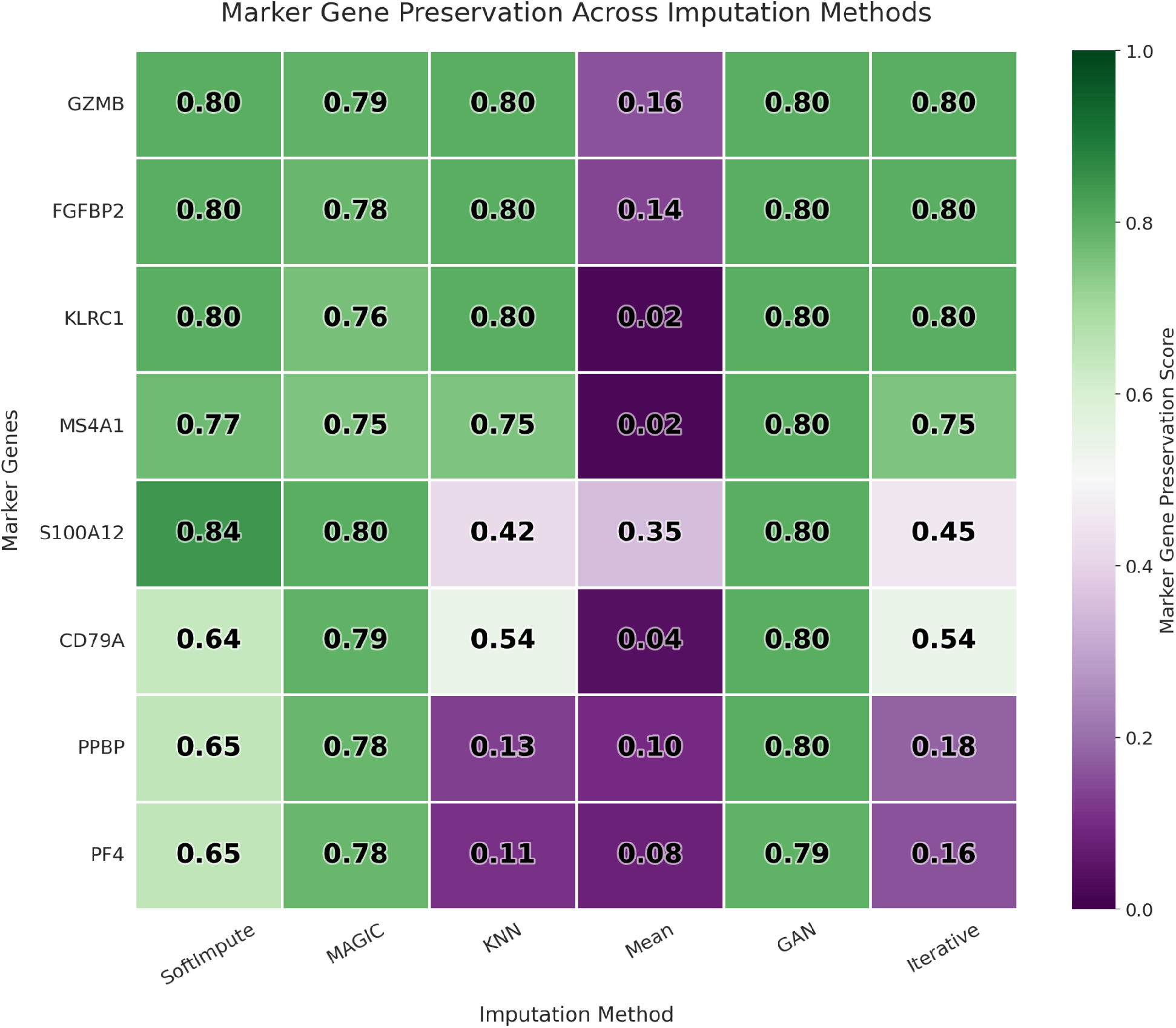
Marker gene preservation quantified according to the color bar across the Imputation methods applied. The marker genes targeted on are given on the left.

In terms of the overall cell clustering consistency, we calculated the Adjusted Rand Index (ARI) relative to the ground truth and summarized the results in Figure 2. SoftImpute demonstrated the strongest and most stable performance, maintaining the highest ARI values across all simulated dropout levels, including 30%. GAN produced clustering results comparable to SoftImpute at 10% dropout, but its performance declined more sharply as missingness increased. MAGIC showed the weakest clustering recovery, whereas Mean and Iterative imputers yielded moderate but consistent values.

**Figure 2:**
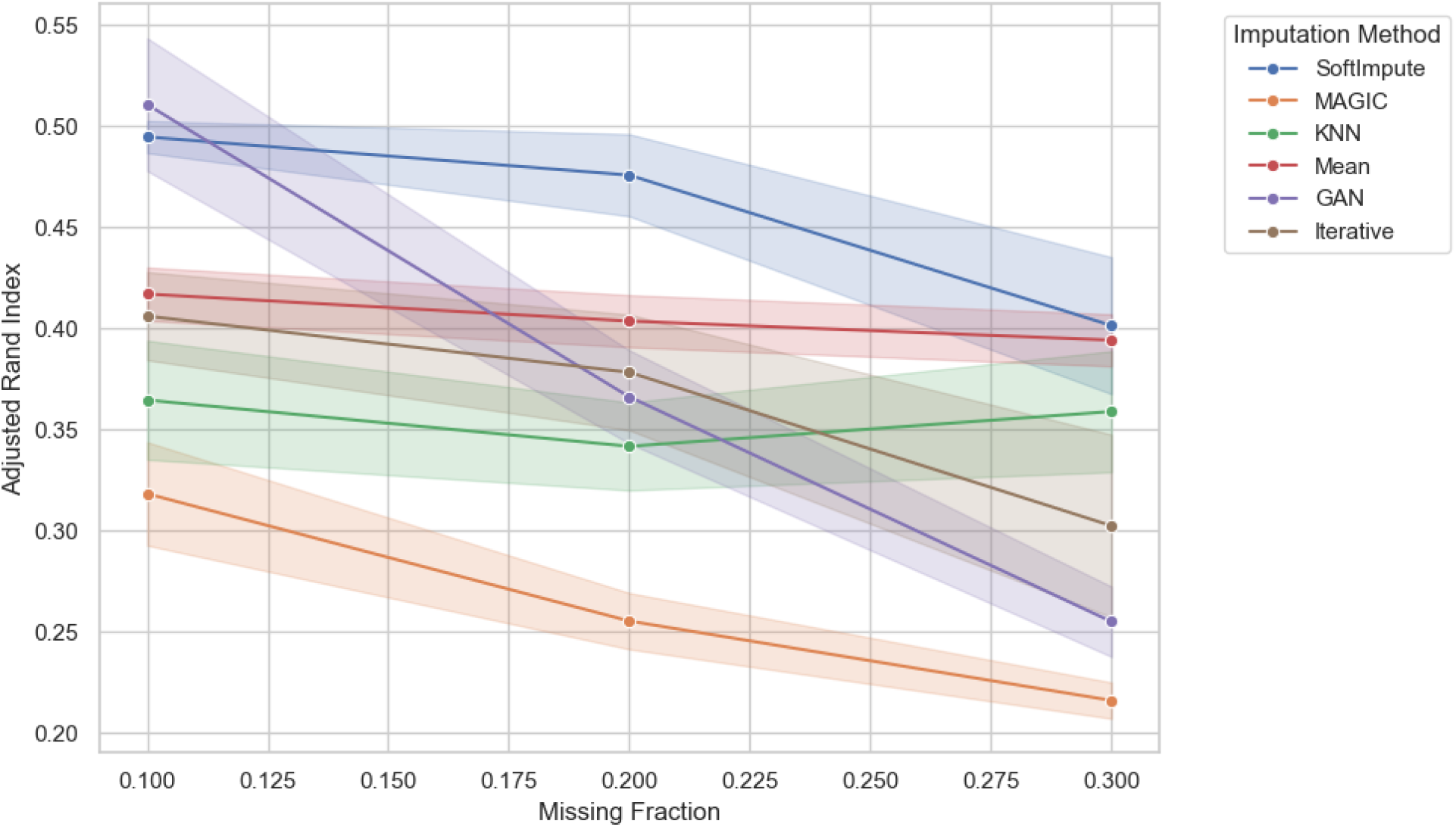
ARI across missing fractions on the x-axis and imputation methods as described in the legend.

On the gene-wise correlation front and the level of individual gene expression recovery, GAN outperformed all other methods to which it is compared in Figure 3. The correlation with the ground truth reached ∼0.95 at 10% dropout and remained the highest among all methods at 30% dropout. SoftImpute and KNN also achieved relatively strong correlations, although consistently lower than GAN. The Iterative method declined steadily with greater dropout, while MAGIC and Mean imputation schemes showed poor performance, averaging around 0.4 regardless of the dropout level.

**Figure 3:**
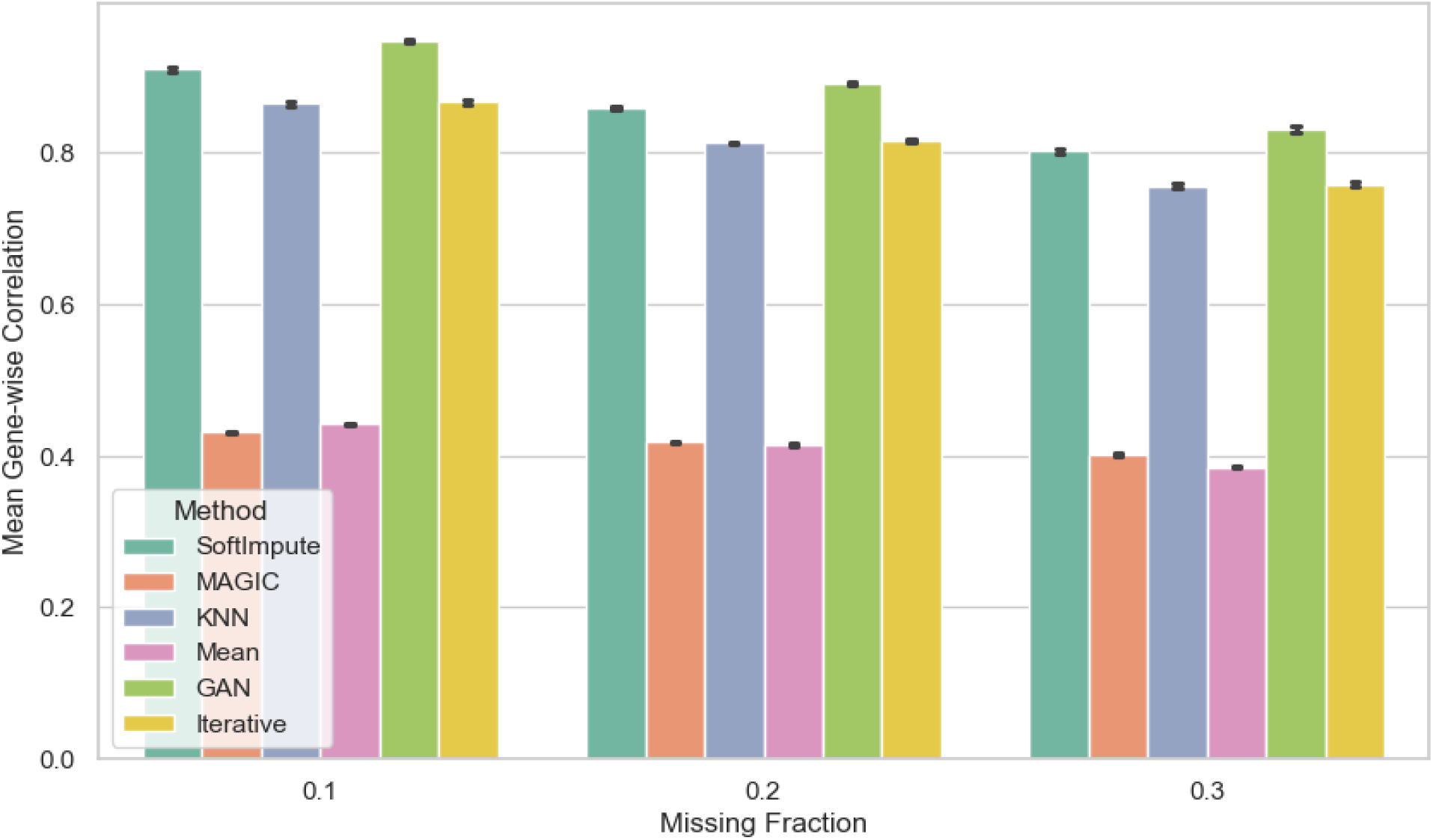
Gene-wise correlation across missing fractions in the x-axis and imputation methods as denoted in the legends.

Finally, we evaluated the structural separation of clusters in reduced-dimensional space using silhouette scores in Figure 4. MAGIC produced the highest silhouette scores across the dropout levels, but this result was inconsistent with its poor ARI and correlation performance, suggesting that the method may over-smooth expression profiles and artificially exaggerate separation. SoftImpute, GAN, and Iterative methods yielded lower or negative silhouette values, reflecting more conservative reconstructions. The Mean and KNN showed intermediate but variable results.

**Figure 4:**
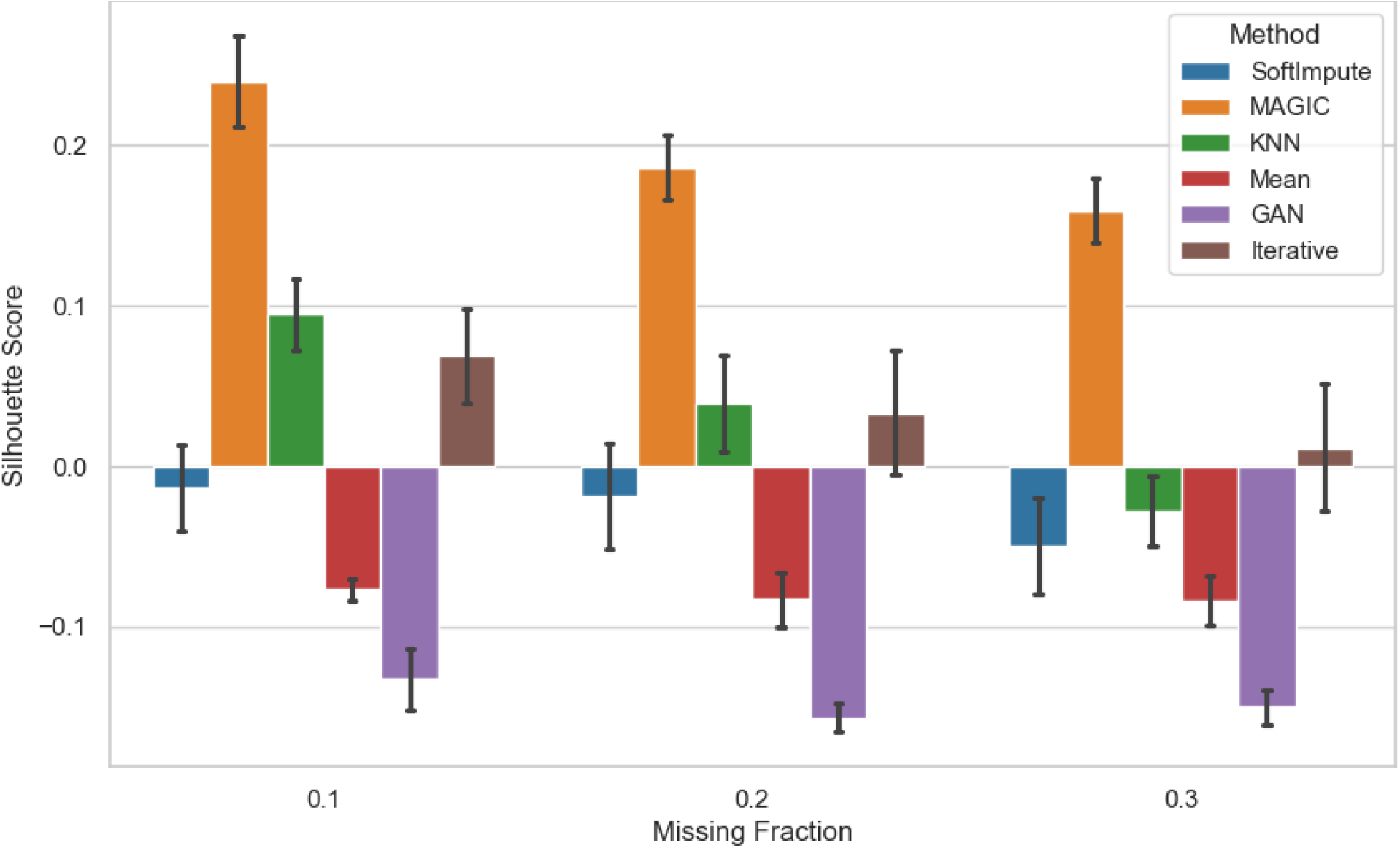
The Silhouette scores across missing fractions on the x-axis and imputation methods as denoted in the legends. The error bars denote the variation over the ten distinct calculations for each case.

## Conclusions

Using the PBMC AMI dataset as the ground truth, this study benchmarked six imputation techniques for AMI-related single-cell RNA sequencing data across three levels of fake missingness. Our findings demonstrate that imputation choice is not a neutral preprocessing step; rather, it actively influences which biological signals are recovered and which are lost, with clear implications for the interpretation of post-AMI immune dynamics.No single method dominated across all metrics: GAN and SoftImpute emerged as the strongest combination, with GAN achieving a global transcriptional recovery score above 0.80 (making it well-suited for capturing overall transcriptional trends and regulatory network inference), while SoftImpute consistently delivered the best clustering fidelity, maintaining ARI values above 0.40 even at 30% dropout, making it the preferred choice when preserving cell-type identity is the priority. On the other hand, simpler approaches like Mean and KNN imputation performed poorly across nearly all benchmarks, which indicates that their over-smoothing behavior risks masking or distorting the immune signatures, including platelet, B-cell, NK-cell, and monocyte markers assessed against the ground truth that are relevant to understanding post-AMI inflammation and repair. The MAGIC result of high silhouette scores alongside weak ARI and correlation performance shows that structural clarity in low-dimensional embeddings does not guarantee true biological recovery and can instead reflect artificial signal inflation.

These results have implications for cardiovascular single-cell research in general, in addition to their immediate applicability. The timing and makeup of immune cell recruitment can determine whether tissue repair proceeds favourably or maladaptively in AMI; selecting an imputation method that obscures rare or transitional immune populations or inflates apparent structure without true signal could significantly skew conclusions about which cell states or marker genes drive disease progression. This directly affects the identification of therapeutic targets and biomarkers in AMI, as downstream experimental validation may be misguided by a false sense of confidence in artificially “cleaned” data. More broadly, our findings suggest that rather than being chosen by default due to computational ease or popularity, imputation should be carefully chosen based on the particular downstream question, such as transcriptome-wide trend detection, marker gene validation, or fine-grained cell-type resolution. The benchmarking framework developed here provides a template that can be applied to other acute inflammatory or ischaemic conditions, helping to ensure that the biological insights derived from imputed data are reliable rather than artefacts of the imputation process itself, as single-cell approaches become more and more important in precision cardiovascular medicine.

A few limitations of the current work should be noted. The artificial missingness was simulated under a missing completely at random (MCAR) assumption, which may not accurately represent the more structured, expression-dependent dropout patterns frequently seen in actual scRNA-seq experiments. Our benchmarking was carried out on a single dataset derived from a single patient cohort. Therefore, without additional validation, the relative efficacy of the imputation techniques described here may not be directly applicable to different tissue types, disease contexts, or sequencing platforms, even though they are resilient across multiple dropout levels and repeated masking runs. Furthermore, our investigation concentrated on a specific collection of marker genes and common clustering/correlation metrics; alternative biologically significant readouts, including trajectory inference or cell-cell communication analysis, were not evaluated and might react differently to imputation selection. Future research expanding this benchmarking methodology to other AMI populations, more missingness mechanisms (such as Missing Not At Random, Missing At Random), and a wider range of downstream analyses would support the findings’ generalisability and bolster their translational significance. Marker gene evaluation was limited to genes present within the ground truth reference’s highly-variable-gene feature set, which meant several canonical markers (e.g., T-cell markers CD3D/E/G, CD4, CD8A/B, and CD14 for classical monocytes) fell outside the compared feature space and could not be assessed in this study.

## Declarations

### Ethics approval and consent to participate

Not applicable

### Consent for publication

Not applicable

### Availability of data and materials

The single-cell RNA sequencing dataset analysed in this study was obtained from the NCBI Gene Expression Omnibus (GEO) under accession ID GSE269269 (*Single-Cell RNA Sequencing of PBMCs from Acute Myocardial Infarction*; https://www.ncbi.nlm.nih.gov/geo/query/acc.cgi?acc=GSE269269)

All analyses were conducted in Python 3.10 on a Linux environment with conda-based environment management. The code is publicly available on GitHub under the link https://github.com/RWTH-CompBioTech/Benchmarking-Imputation-Methods-in-Single-cell-RNA-Seq-of-PBMCs-from-Acute-Myocardial-Infarction

## Acknowledgements

PR is grateful to Prof. Sharmistha Majumdar, Associate Professor, Department of Biological Engineering, Indian Institute of Technology Gandhinagar, for supporting this work. The authors gratefully acknowledge the computing time provided to them at the NHR Center NHR4CES at RWTH Aachen University. This is funded by the Federal Ministry of Education and Research, and the state governments participating on the basis of the resolutions of the GWK for national high performance computing at universities.

## Competing interests

The authors declare that they have no competing interests

## Funding

PR was supported by the DAAD KOSPIE Scholarship to conduct her master’s thesis research at RWTH Aachen University in collaboration with IIT Gandhinagar. This support facilitated the development of the present research project in collaboration with MF.

## Authors’ contributions

PR designed and executed the research, performed the analyses, and prepared the initial draft. MF approved the research proposal, provided guidance throughout the study, and finalized the manuscript. All authors read and approved the final manuscript.

## Notes

### Competing Interest Statement

The authors have declared no competing interest.

